## Supplementary material for "Single Cell Transcriptomic Landscape of Diabetic Foot Ulcers": Supplmentary Information

### Equal First authors

* Co-last and corresponding authors

***Corresponding authors***:

Aristidis Veves, MD, DSc.
Beth Israel Deaconess Medical Center,
Palmer 321A, 1 Deaconess Rd,
Boston, MA 02215.


Manoj K. Bhasin, MS, PhD.
Aflac Cancer and Blood Disorders Center
Children Healthcare of Atlanta
Woodruff Memorial Research Building, Room 4107
101 Woodruff Circle, 4th Floor East
Emory School of Medicine
Atlanta, GA 30322.


#### Supplementary figures

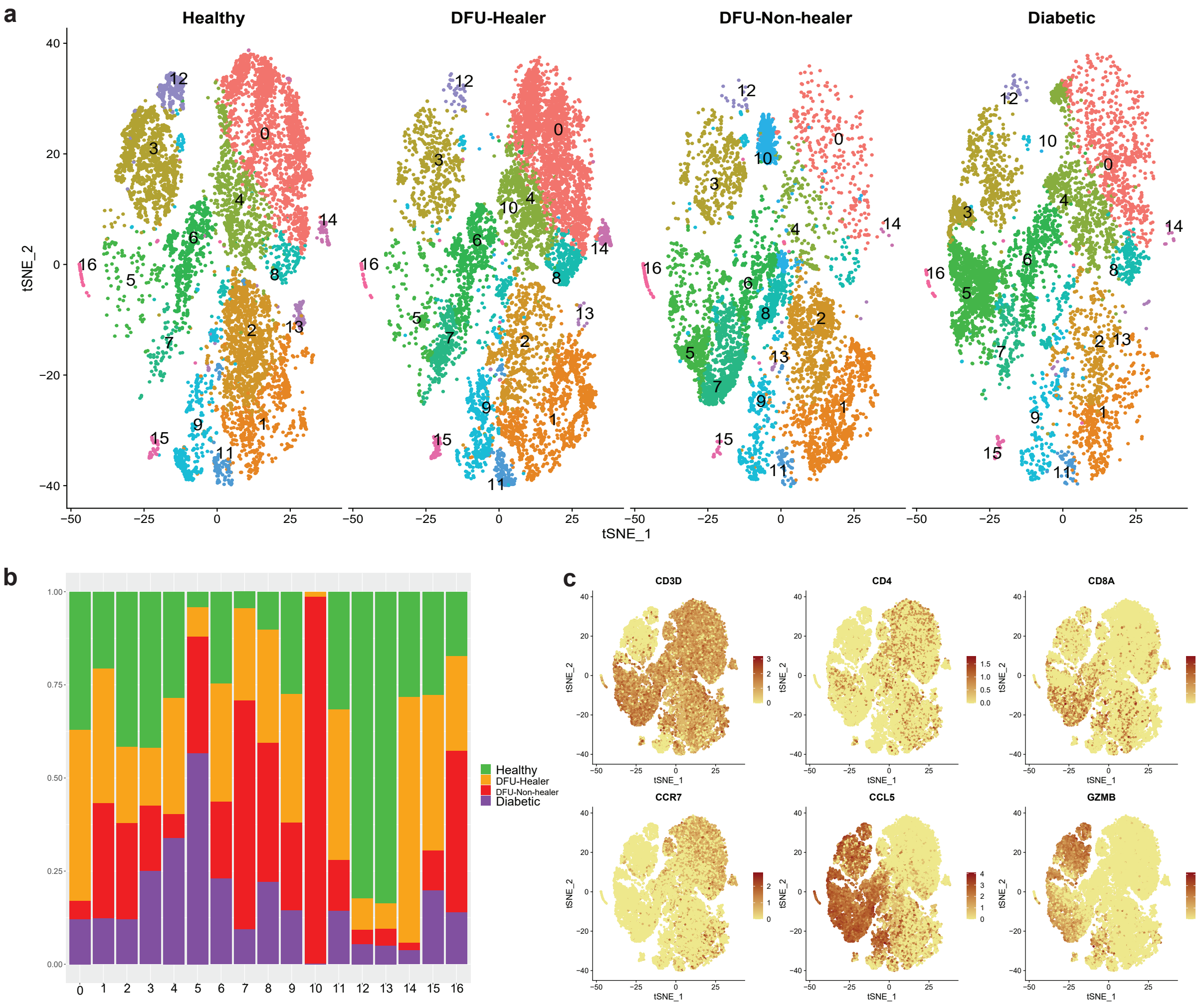


**Figure S1. *Comparative analysis of T, NKT, and NK cell subpopulations in different clinical groups.***

**(a)** t-distributed Stochastic Neighbor Embedding (t-SNE) analysis of T-lymphocytes, Natural Killer (NK) cells and NKT cells, **(b)** The subcluster wise proportion of different cell subtypes across clinical groups (i.e., Green: Healthy non-DM, Orange: DFU-Healers, Red: DFU-Non-Healers, Purple: non-DFU DM). **(c)** Feature maps showing expression of gene markers for T cells (*CD3D*$^{+}$), T-helper (*CD4*$^{+}$), T-cytotoxic (*CD8A*$^{+}$), Naïve/Central Memory T-cell (*CCR7*$^{+}$), effector CD8$^{+}$ T cells (*CCL5*$^{+}$), and NK (*GZMB*$^{+}$ *and CD3D*$^{-}$).

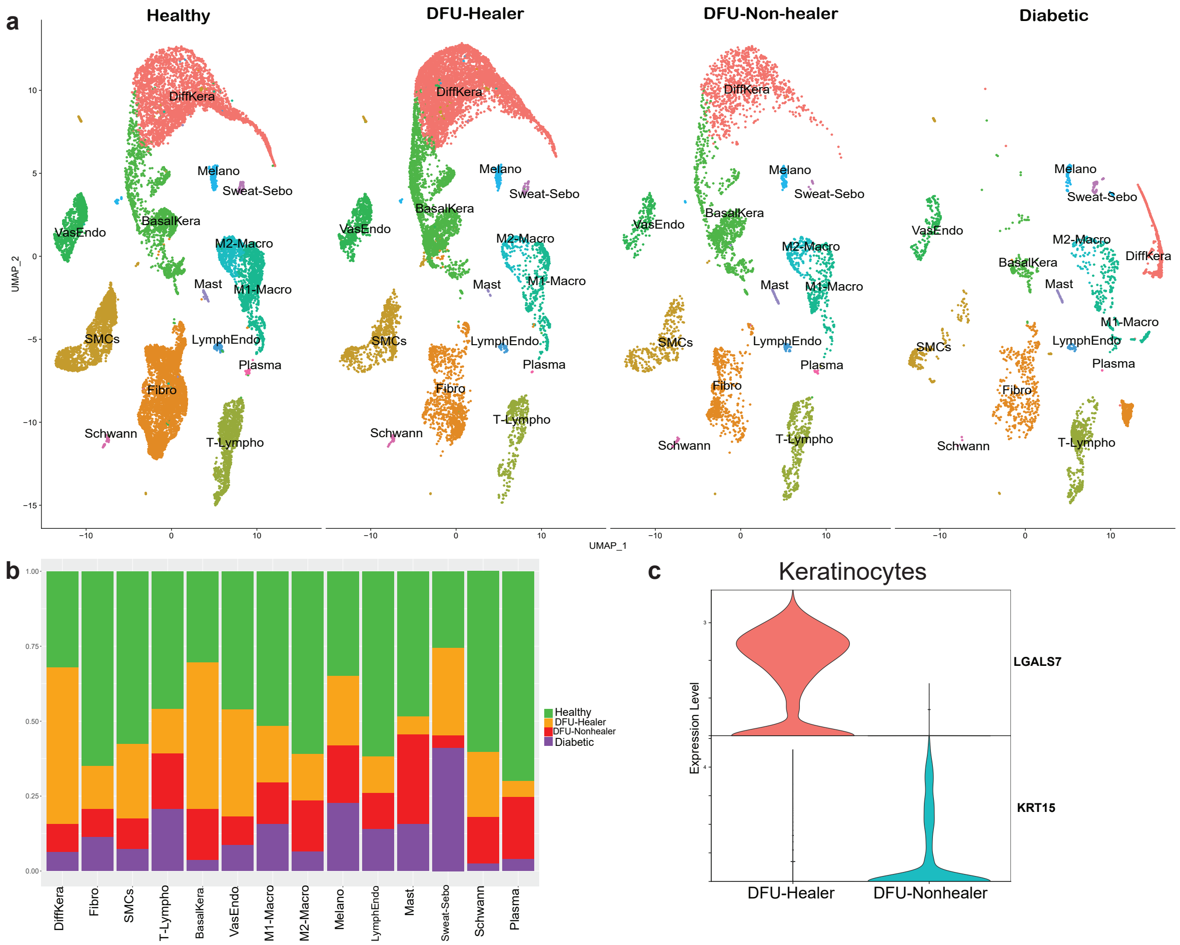


**Figure S2.** ***Comparative analysis of transcriptome profiles of forearm biopsies in the different clinical groups***.

**(a)** UMAP dimensionality reduction embedding of forearm skin cells from DFU-Healers, DFU-Non-healers, Healthy subjects, and non-DFU DM patients. **(b)** Stacked bar plots showing the proportions of different cell types across the different clinical groups (Green: Healthy non-DM, Orange: DFU-Healers, Red: DFU-Non-Healers, Purple: non-DFU DM). **(c)** Violin plots showing expression levels of top differentially expressed gene *LGALS7* in DFU-Healers and *KRT15* in DFU-Non-healers.

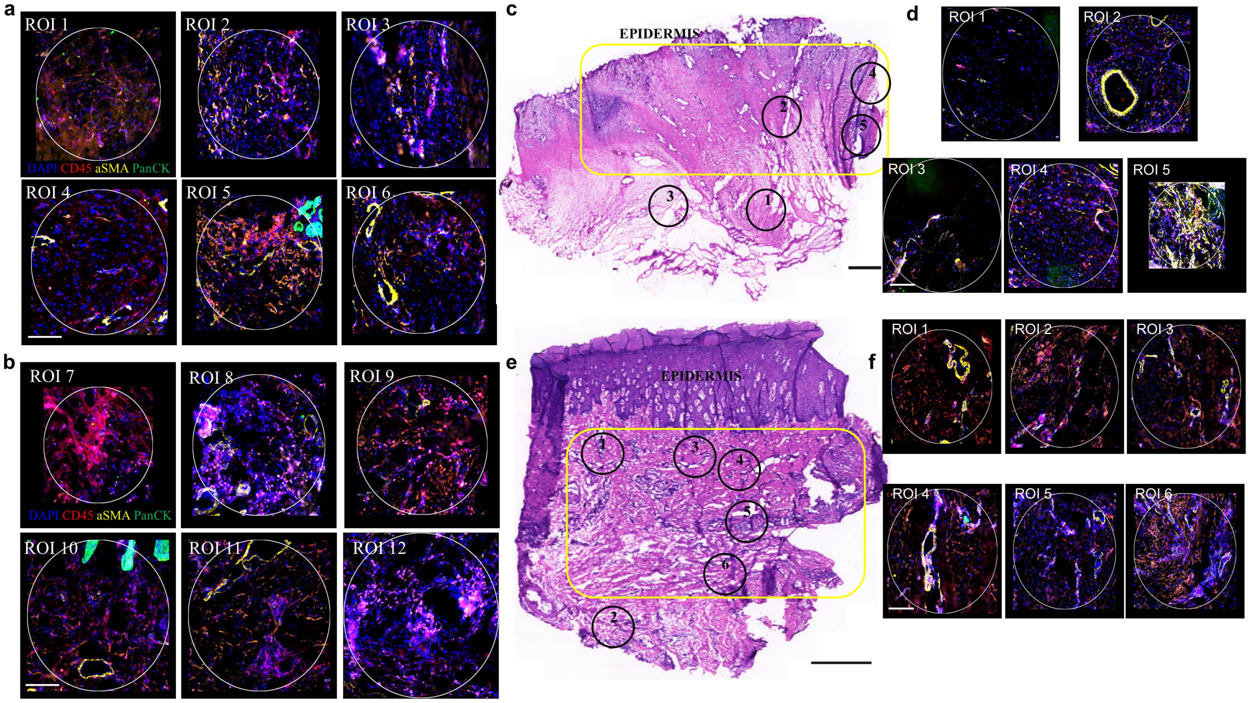


**Figure S3. Regions of interest selection** (**a, b**) Micrographs presenting the ROIs selected for spatial transcriptomics analysis from a non-healing (**a**) and healing (**b**) DFU labeled by immunofluorescence as shown of Figure 6. (**c,e**) Representative H&E stained sections from a non-healing (**c**) and a healing (**e**) DFU. Yellow box demarcates the ulcer area and numbered circles the ROIs selected for sequencing. (**d,f**) The immunofluorescence staining of ROIs for markers CD45 (red), aSMA (yellow) and panCytokeratin (green). DAPI was used for nuclear counterstain. Scale bars are 100 μm in (**a,b,d,f**) and 1 mm in (**c,e**).

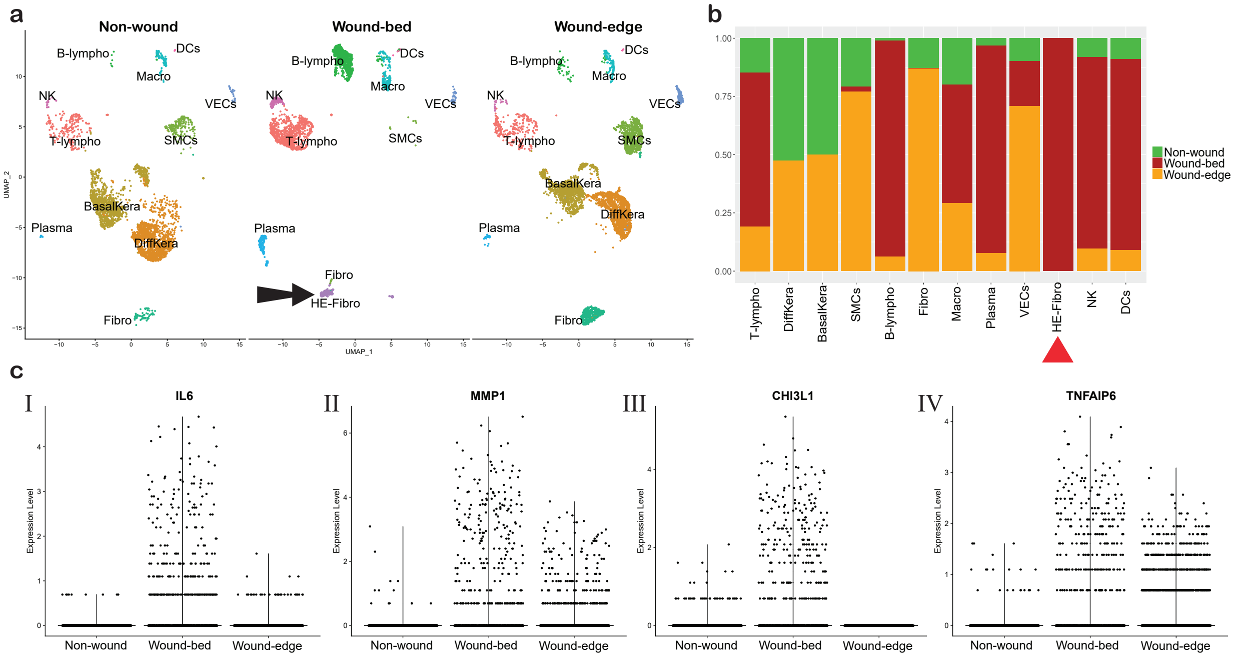


**Figure S4. Healing Enriched Fibroblasts (HE-Fibro) are exclusively localized to wound bed. (a)** UMAP showing distribution of different cell types in three samples from the non-wound site, wound bed and edge of the wound of one patient. Black arrow indicates the HE-Fibro cell cluster. **(b)** Bar plot showing relative proportions of cells from the 3 samples in the different cell types. Red triangle indicates the HE-Fibro cell cluster exclusively present in the wound bed. **(c)** Expression levels of (I) *IL6*, (II) *MMP1*, (III) *CHI3L1*, (IV) *TNFAIP6*, which were found to be overexpressed by the HE-Fibro cells in our study, showed higher expression in the wound bed as compared to the non-wound site.

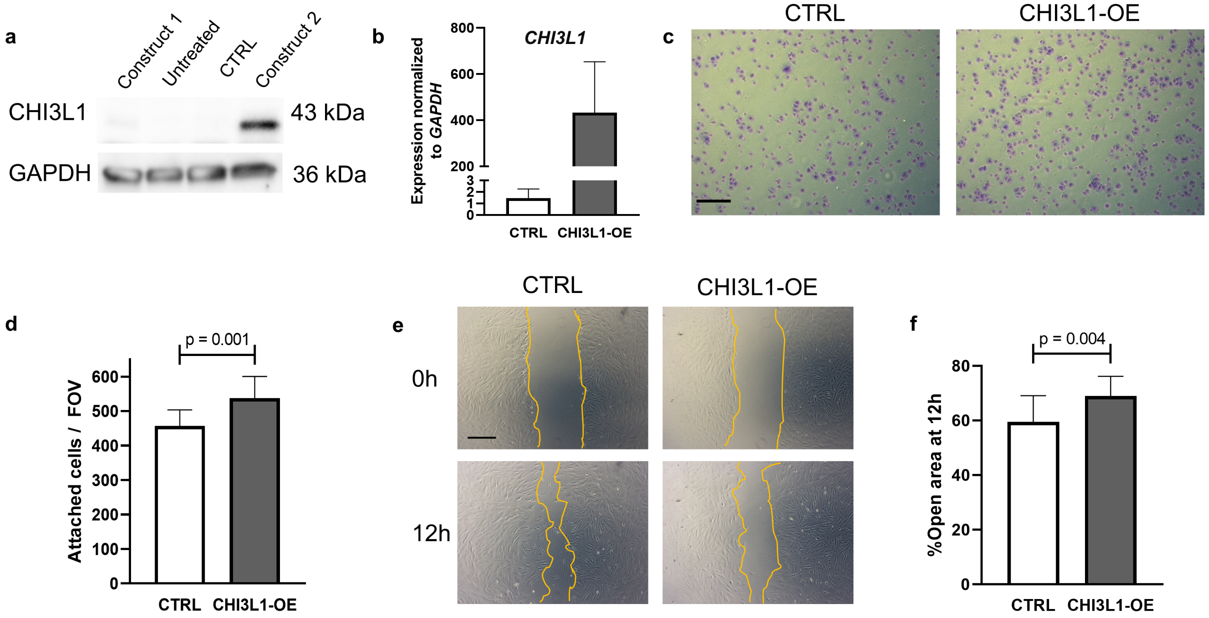


**Figure S5 CHI3L1 overexpression in dermal fibroblasts** (a) Western blot analysis of two *CHI3L1* constructs, 1 and 2, that overexpress CHI3L1, untreated cells and control construct (CTRL) expressing cells. GAPDH was used as a loading control. (b) RT-qPCR of CTRL and Construct 2 (CHI3L1-OE) expressing cells. (c, d) Representative images (c) and quantitation (d) of crystal violet stained CTRL and CHI3L1-OE cells that attached after 1 h on fibronectin coated wells. (e, f) Representative brightfield images (e) and quantitation (f) of scratch wounds at 0 and 12 h. Yellow lines denote migrated cells. Data represents mean±SD of N=12-16 observations from three independent experiments. P values were calculated by two-tailed unpaired t test. Scale bars are 100 μm. FOV: field of view.

**Supplementary Table 1: Clinical characteristics of patients across groups.**

| | **Healthy Controls** | **DM without DFU** | **DM-DFU** |
|---|---|---|---|
| Number (Males) | 10 (4) | 6 (2) | 11 (4) |
| Age (years) | 59 ± 13 | 64 ±12 | 55 ± 16 |
| DM duration (years) | -- | 10 ± 6 | 16 ± 15 |
| BMI | 28.3 ± 3.8 | 32.1 ± 8.8 | 38.6 ± 12.3 |
| HbA1c* | 5.5 ± 0.7 | 8.4 ± 1.7 | 8.8 ± 2.6 |
| Creatinine | 0.9 ± 0.2 | 1.1 ± 0.4 | 1.4 ± 1.0 |

*: Healthy Controls vs DM without DFU and DM with DFU; $p<0.05$, ANOVA with Fisher's LSD *post-hoc*.

**Supplementary Table 2: List of Top 10 genes overexpressed in the cell types clusters.** Table showing top ten marker genes for each cell type cluster with average fold-change and adjusted p value. The fold change was calculated by comparing the expression profiles of target cell type cluster with other cell types.

| Cell type | Gene | Avg. log FC | Adj. p value |
|---|---|---|---|
| Fibro | *APOD* | 2.97 | 0 |
| | *CFD* | 2.79 | 0 |
| | *DCN* | 2.77 | 0 |
| | *COL1A1* | 2.59 | 0 |
| | *COL1A2* | 2.56 | 0 |
| | *SFRP2* | 2.34 | 0 |
| | *COL3A1* | 2.32 | 0 |
| | *PTGDS* | 2.29 | 0 |
| | *CXCL14* | 2.28 | 0 |
| | *COMP* | 2.21 | 0 |
| SMCs | *TAGLN* | 3.07 | 0 |
| | *ACTA2* | 2.98 | 0 |
| | *TPM2* | 2.33 | 0 |
| | *MYL9* | 2.27 | 0 |
| | *RGS5* | 2.27 | 0 |
| | *C11orf96* | 2.20 | 0 |
| | *CALD1* | 1.91 | 0 |
| | *IGFBP7* | 1.75 | 0 |
| | *MYH11* | 1.71 | 0 |
| | *TPM1* | 1.66 | 0 |
| CD14-Mono | *S100A9* | 3.72 | 0 |
| | *LYZ* | 3.35 | 0 |
| | *S100A8* | 3.08 | 0 |
| | *FCN1* | 2.48 | 0 |
| | *CTSS* | 2.48 | 0 |
| | *MNDA* | 2.11 | 0 |
| | *S100A12* | 1.95 | 0 |
| | *VCAN* | 1.91 | 0 |
| | *TYROBP* | 1.83 | 0 |
| | AIF1 | 1.76 | 0 |
| VasEndo | IFI27 | 2.21 | 0 |
| | ACKR1 | 2.12 | 0 |
| | CLDN5 | 1.88 | 0 |
| | AQP1 | 1.84 | 0 |
| | GNG11 | 1.84 | 0 |

| | | | |
|---|---|---|---|
| | SPARCL1 | 1.81 | 0 |
| | PECAM1 | 1.79 | 0 |
| | VWF | 1.75 | 0 |
| | PLVAP | 1.75 | 0 |
| | RAMP2 | 1.71 | 0 |
| T-lympho | LTB | 1.61 | 0 |
| | IL7R | 1.45 | 0 |
| | IL32 | 1.30 | 0 |
| | TRAC | 1.15 | 0 |
| | CD69 | 1.12 | 0 |
| | CD3D | 1.11 | 0 |
| | TRBC2 | 1.00 | 0 |
| | CD52 | 0.95 | 0 |
| | CXCR4 | 0.92 | 0 |
| | TRBC1 | 0.91 | 0 |
| DiffKera | KRT10 | 3.78 | 0 |
| | KRT2 | 3.64 | 0 |
| | KRT1 | 3.49 | 0 |
| | DMKN | 3.36 | 0 |
| | KRTDAP | 3.32 | 0 |
| | CALML5 | 2.64 | 0 |
| | PERP | 2.51 | 0 |
| | LGALS7B | 2.48 | 0 |
| | DSP | 2.41 | 0 |
| | SBSN | 2.41 | 0 |
| BasalKera | KRT14 | 4.28 | 0 |
| | KRT5 | 3.62 | 0 |
| | KRT16 | 3.12 | 0 |
| | KRT6A | 2.76 | 0 |
| | S100A2 | 2.56 | 0 |
| | KRT17 | 2.54 | 0 |
| | KRT6B | 2.35 | 0 |
| | SFN | 1.74 | 0 |
| | COL17A1 | 1.64 | 0 |
| | KRT15 | 1.47 | 0 |
| HE-Fibro | PLA2G2A | 2.92 | 0 |
| | MMP1 | 2.53 | 0 |
| | CHI3L1 | 2.48 | 0 |
| | TIMP1 | 2.11 | 0 |
| | SFRP4 | 2.01 | 0 |
| | FTH1 | 1.81 | 0 |
| | FN1 | 1.80 | 0 |
| | CHI3L2 | 1.80 | 0 |

| | | | |
|---|---|---|---|
| | MT2A | 1.77 | 0 |
| | LUM | 1.74 | 0 |
| B-lympho | IGHM | 2.61 | 0 |
| | IGKC | 1.97 | 0 |
| | CD37 | 1.80 | 0 |
| | MS4A1 | 1.78 | 0 |
| | CD79A | 1.74 | 0 |
| | LTB | 1.71 | 0 |
| | CD79B | 1.53 | 0 |
| | CD52 | 1.40 | 0 |
| | CD74 | 1.37 | 0 |
| | IGLC3 | 1.33 | 0 |
| M1-Macro | HLA-DRA | 2.79 | 0 |
| | IL1B | 2.77 | 0 |
| | HLA-DPB1 | 2.72 | 0 |
| | HLA-DRB1 | 2.67 | 0 |
| | HLA-DPA1 | 2.62 | 0 |
| | CXCL8 | 2.53 | 0 |
| | HLA-DQA1 | 2.50 | 0 |
| | HLA-DQB1 | 2.45 | 0 |
| | CD74 | 2.29 | 0 |
| | G0S2 | 2.16 | 0 |
| NK | GNLY | 3.43 | 0 |
| | NKG7 | 2.82 | 0 |
| | CCL5 | 2.22 | 0 |
| | GZMB | 2.01 | 0 |
| | GZMA | 1.94 | 0 |
| | CST7 | 1.76 | 0 |
| | FGFBP2 | 1.74 | 0 |
| | PRF1 | 1.73 | 0 |
| | CCL4 | 1.57 | 0 |
| | GZMH | 1.55 | 0 |
| NKT | CCL5 | 2.09 | 0 |
| | NKG7 | 1.82 | 0 |
| | GZMA | 1.42 | 0 |
| | IL32 | 1.42 | 0 |
| | GZMH | 1.34 | 0 |
| | GNLY | 1.27 | 0 |
| | CD3D | 1.17 | 0 |
| | CST7 | 1.14 | 0 |
| | GZMK | 1.12 | 0 |
| | IL7R | 1.11 | 0 |
| M2-Macro | RNASE1 | 3.25 | 0 |

| | | | |
|---|---|---|---|
| | C1QA | 2.97 | 0 |
| | SELENOP | 2.51 | 0 |
| | C1QB | 2.46 | 0 |
| | C1QC | 2.21 | 0 |
| | CCL3 | 2.06 | 0 |
| | CD74 | 2.05 | 0 |
| | FTL | 2.02 | 0 |
| | CCL18 | 2.00 | 0 |
| | FOLR2 | 1.92 | 0 |
| CD16-Mono | LST1 | 2.65 | 0 |
| | FCGR3A | 2.61 | 0 |
| | AIF1 | 2.36 | 0 |
| | FCER1G | 2.16 | 0 |
| | COTL1 | 2.08 | 0 |
| | CTSS | 2.01 | 0 |
| | CDKN1C | 1.91 | 0 |
| | MS4A7 | 1.85 | 0 |
| | TYROBP | 1.76 | 0 |
| | SERPINA1 | 1.65 | 0 |
| Melano/Scwann | DCT | 3.65 | 0 |
| | TYRP1 | 3.39 | 0 |
| | PMEL | 2.98 | 0 |
| | MLANA | 2.60 | 0 |
| | GPM6B | 1.79 | 0 |
| | PLP1 | 1.75 | 0 |
| | QPCT | 1.70 | 0 |
| | PMP22 | 1.62 | 0 |
| | MFSD12 | 1.53 | 0 |
| | CYB561A3 | 1.48 | 0 |
| Mast | TPSB2 | 3.45 | 0 |
| | TPSAB1 | 3.25 | 0 |
| | CTSG | 2.70 | 0 |
| | HPGD | 2.62 | 0 |
| | CPA3 | 1.65 | 0 |
| | HPGDS | 1.60 | 0 |
| | GATA2 | 1.45 | 0 |
| | MS4A2 | 1.32 | 0 |
| | LGALS3 | 1.29 | 0 |
| | ANXA1 | 1.25 | 0 |
| Sweat/Sebo | DCD | 6.46 | 0 |
| | SCGB2A2 | 4.83 | 0 |
| | MUCL1 | 4.41 | 0 |
| | SCGB1B2P | 3.85 | 0 |

| | | | |
|---|---|---|---|
| | SCGB1D2 | 3.56 | 0 |
| | PIP | 3.47 | 0 |
| | AQP5 | 2.75 | 0 |
| | KRT19 | 2.74 | 0 |
| | AZGP1 | 2.54 | 0 |
| | SLC12A2 | 2.37 | 0 |
| LymphEndo | CCL21 | 4.57 | 0 |
| | TFF3 | 3.01 | 0 |
| | MMRN1 | 2.27 | 0 |
| | CLDN5 | 1.93 | 0 |
| | GNG11 | 1.93 | 0 |
| | TFPI | 1.75 | 0 |
| | PPFIBP1 | 1.69 | 0 |
| | CAVIN2 | 1.58 | 0 |
| | LYVE1 | 1.56 | 0 |
| | LMO2 | 1.49 | 0 |
| Erytho | HBB | 8.06 | 0 |
| | HBA2 | 6.85 | 0 |
| | HBA1 | 6.50 | 0 |
| | HBD | 2.89 | 0 |
| | ALAS2 | 2.62 | 0 |
| | SNCA | 2.07 | 0 |
| | AHSP | 2.03 | 0 |
| | HBM | 1.78 | 0 |
| | CA1 | 1.58 | 0 |
| | UBB | 1.48 | 0 |
| Plasma | IGKC | 6.71 | 0 |
| | IGLC2 | 6.67 | 0 |
| | IGHA1 | 6.08 | 0 |
| | IGHG1 | 5.94 | 0 |
| | IGLC3 | 5.70 | 0 |
| | IGHG2 | 5.02 | 0 |
| | IGHG3 | 4.90 | 0 |
| | JCHAIN | 4.62 | 0 |
| | IGHG4 | 3.92 | 0 |
| | MZB1 | 2.90 | 0 |
| DCs | GZMB | 2.60 | 0 |
| | JCHAIN | 1.90 | 0 |
| | IRF7 | 1.88 | 0 |
| | IRF8 | 1.87 | 0 |
| | PLAC8 | 1.74 | 0 |
| | PLD4 | 1.68 | 0 |
| | ALOX5AP | 1.63 | 0 |

| | | | |
|---|---|---|---|
| | CCDC50 | 1.76 | 2.58E-210 |
| | ITM2C | 1.69 | 2.24E-196 |
| | PTGDS | 1.70 | 2.23E-33 |

**Supplementary table 3:** Distribution of different cell types across anatomical locations.

| Cell type | Average % of cells ± SE | | |
|---|---|---|---|
| | Foot | forearm | PBMC |
| Fibro | 73.54 ± 0.50 | 26.46 ± 0.54 | |
| SMCs | 86.65 ± 0.61 | 13.35 ± 0.22 | |
| CD14-Mono | 1.53 ± 0.04 | | 98.47 ± 1.06 |
| VasEndo | 82.37 ± 0.33 | 17.63 ± 0.29 | |
| T-lympho | 31.27 ± 0.27 | 12.42 ± 0.35 | 56.31 ± 1.21 |
| DiffKera | 17.26 ± 0.69 | 73.28 ± 1.73 | |
| BasalKera | 71.35 ± 0.48 | 28.65 ± 0.72 | |
| HE-fibro | 99.94 ± 1.58 | 0.06 ± 0.00 | |
| B-Lympho | 15.51 ± 0.27 | | 84.49 ± 1.63 |
| M1-Macro | 55.23 ± 0.55 | 36.75 ± 0.65 | 8.02 ± 0.11 |
| Nk | 12.08 ± 0.15 | 2.88 ± 0.06 | 85.04 ± 1.11 |
| NKT | 3.78 ± 0.04 | 2.01 ± 0.07 | 94.21 ± 3.62 |
| M2-Macro | 71.28 ± 1.27 | 28.62 ± 1.09 | 0.10 ± 0.01 |
| CD16-Mono | 0.28 ± 0.01 | 0.09 ± 0.00 | 99.63 ± 1.11 |
| Melano/Schwann | 58.01 ± 0.76 | 41.99 ± 0.81 | |
| Mast | 82.51 ± 1.49 | 17.34 ± 0.74 | 0.16 ± 0.01 |
| Sweat/Seba | 85.96 ± 0.74 | 14.04 ± 0.41 | |
| LymphEndo | 75.42 ± 0.39 | 24.58 ± 0.53 | |
| Erythro | 1.72 ± 0.06 | | 98.28 ± 5.82 |
| Plasma | 66.01 ± 1.89 | 19.38 ± 1.18 | 14.61 ± 0.46 |
| DCs | 1.92 ± 0.04 | 4.49 ± 0.38 | 93.59 ± 1.99 |

Table showing average percentage of each cell type in three samples types, namely foot, forearm and PBMCs. The cell type percentage were calculated as percent of total cells of the specific cell type from the three anatomical sites of collection. SE was calculated as square root of standard deviation between the number of cells present in each patient sample.

**Supplementary Table 4: List of Top 10 genes overexpressed in the foot cell types.** Table showing top ten marker genes for each cell type with average fold-change and adjusted p value. The fold change was calculated by comparing the expression profiles of target cell type with rest of the cell types.

| Cell Type | Gene | Average log fold change | adjusted p value |
|---|---|---|---|
| SMC1 | *ACTA2* | 2.76 | 0 |
| | *TAGLN* | 2.72 | 0 |
| | *RGS5* | 2.28 | 0 |
| | *TPM2* | 2.09 | 0 |
| | *MYL9* | 2.01 | 0 |
| | *C11orf96* | 2.00 | 0 |
| | *MYH11* | 1.75 | 0 |
| | *NDUFA4L2* | 1.57 | 0 |
| | *CALD1* | 1.54 | 0 |
| | *NR2F2* | 1.48 | 0 |
| VasEndo | *IFI27* | 2.06 | 0 |
| | *ACKR1* | 2.06 | 0 |
| | *PECAM1* | 1.92 | 0 |
| | *CLDN5* | 1.83 | 0 |
| | *PLVAP* | 1.77 | 0 |
| | *VWF* | 1.76 | 0 |
| | *SOX18* | 1.72 | 0 |
| | *GNG11* | 1.69 | 0 |
| | *RAMP2* | 1.68 | 0 |
| | *AQP1* | 1.66 | 0 |
| Fibro | *CFD* | 3.03 | 0 |
| | *APOD* | 2.96 | 0 |
| | *PLA2G2A* | 2.54 | 0 |
| | *DCN* | 2.46 | 0 |
| | *CXCL14* | 2.41 | 0 |
| | *PTGDS* | 2.33 | 0 |
| | *SFRP2* | 2.25 | 0 |
| | *COMP* | 2.01 | 0 |
| | *FBLN1* | 1.84 | 0 |
| | *GSN* | 1.77 | 0 |
| T-Lympho | *CD69* | 1.71 | 0 |
| | *IL32* | 1.65 | 0 |
| | *CXCR4* | 1.54 | 0 |
| | *CD52* | 1.51 | 0 |
| | *LTB* | 1.51 | 0 |
| | *KLRB1* | 1.46 | 0 |

| | | | |
|---|---|---|---|
| | *IL7R* | 1.34 | 0 |
| | *PTPRC* | 1.24 | 0 |
| | *DUSP2* | 1.21 | 0 |
| | *TRAC* | 1.18 | 0 |
| DiffKera | *KRT1* | 4.04 | 0 |
| | *KRT10* | 3.99 | 0 |
| | *KRT2* | 3.44 | 0 |
| | *DMKN* | 3.25 | 0 |
| | *KRTDAP* | 2.74 | 0 |
| | *CALML5* | 2.55 | 0 |
| | *LGALS7B* | 2.47 | 0 |
| | *LY6D* | 2.41 | 0 |
| | *DSP* | 2.31 | 0 |
| | *PERP* | 2.28 | 0 |
| M1-Macro | *LYZ* | 3.16 | 0 |
| | *IL1B* | 3.01 | 0 |
| | *HLA-DRA* | 2.91 | 0 |
| | *HLA-DPB1* | 2.72 | 0 |
| | *CXCL8* | 2.70 | 0 |
| | *HLA-DRB1* | 2.61 | 0 |
| | *HLA-DPA1* | 2.60 | 0 |
| | *SRGN* | 2.41 | 0 |
| | *HLA-DQA1* | 2.34 | 0 |
| | *CD74* | 2.27 | 0 |
| HE-Fibro | *MMP1* | 2.47 | 0 |
| | *COL1A1* | 2.13 | 0 |
| | *ASPN* | 2.02 | 0 |
| | *POSTN* | 2.01 | 0 |
| | *COL3A1* | 1.92 | 0 |
| | *COL1A2* | 1.85 | 0 |
| | *COL12A1* | 1.77 | 0 |
| | *TNC* | 1.71 | 0 |
| | *LUM* | 1.66 | 0 |
| | *FN1* | 1.61 | 0 |
| BasalKera | *KRT14* | 3.13 | 0 |
| | *KRT16* | 2.99 | 0 |
| | *KRT6A* | 2.72 | 0 |
| | *KRT5* | 2.69 | 0 |
| | *KRT17* | 2.65 | 0 |
| | *KRT6B* | 2.58 | 0 |
| | *KRT6C* | 2.53 | 0 |
| | *S100A2* | 2.51 | 0 |
| | *S100A8* | 2.35 | 0 |

| | | | |
|---|---|---|---|
| | *S100A9* | 2.26 | 0 |
| M2-Macro | *RNASE1* | 3.18 | 0 |
| | *C1QA* | 3.17 | 0 |
| | *C1QB* | 2.70 | 0 |
| | *SELENOP* | 2.49 | 0 |
| | *C1QC* | 2.40 | 0 |
| | *FTL* | 2.30 | 0 |
| | *CD74* | 2.24 | 0 |
| | *CD14* | 2.19 | 0 |
| | *AIF1* | 2.18 | 0 |
| | *HLA-DRA* | 2.18 | 0 |
| Mast | *TPSB2* | 3.54 | 0 |
| | *TPSAB1* | 3.35 | 0 |
| | *CTSG* | 2.79 | 0 |
| | *HPGD* | 2.76 | 0 |
| | *HPGDS* | 1.70 | 0 |
| | *CPA3* | 1.69 | 0 |
| | *GATA2* | 1.42 | 0 |
| | *MS4A2* | 1.39 | 0 |
| | *LGALS3* | 1.37 | 0 |
| | *SERPINB1* | 1.23 | 0 |
| NKT | *GNLY* | 2.80 | 0 |
| | *NKG7* | 2.54 | 0 |
| | *CCL5* | 2.54 | 0 |
| | *CCL4* | 2.49 | 0 |
| | *XCL2* | 1.91 | 0 |
| | *DUSP2* | 1.91 | 0 |
| | *CD69* | 1.90 | 0 |
| | *XCL1* | 1.76 | 0 |
| | *GZMA* | 1.71 | 0 |
| | *CCL3* | 1.68 | 0 |
| B-Lympho | *CD37* | 1.82 | 0 |
| | *LTB* | 1.77 | 0 |
| | *MS4A1* | 1.70 | 0 |
| | *CD74* | 1.70 | 0 |
| | *CD52* | 1.57 | 0 |
| | *HLA-DRA* | 1.54 | 0 |
| | *HLA-DPB1* | 1.40 | 0 |
| | *CD79A* | 1.34 | 0 |
| | *HLA-DQA1* | 1.33 | 0 |
| | *IGHM* | 1.31 | 0 |
| LymphEndo | *CCL21* | 4.64 | 0 |
| | *TFF3* | 2.94 | 0 |

| | | | |
|---|---|---|---|
| | *MMRN1* | 2.22 | 0 |
| | *CLDN5* | 1.77 | 0 |
| | *GNG11* | 1.70 | 0 |
| | *TFPI* | 1.61 | 0 |
| | *PPFIBP1* | 1.60 | 0 |
| | *CAVIN2* | 1.49 | 0 |
| | *PROX1* | 1.46 | 0 |
| | *LMO2* | 1.45 | 0 |
| Melano | *DCT* | 4.08 | 0 |
| | *TYRP1* | 3.94 | 0 |
| | *PMEL* | 3.56 | 0 |
| | *MLANA* | 3.10 | 0 |
| | *QPCT* | 2.21 | 0 |
| | *CYB561A3* | 1.97 | 0 |
| | *MITF* | 1.91 | 0 |
| | *APOE* | 1.86 | 0 |
| | *MFSD12* | 1.75 | 0 |
| | *PLP1* | 1.67 | 0 |
| SMC2 | *CENPF* | 1.92 | 0 |
| | *PTTG1* | 1.87 | 0 |
| | *H2AFZ* | 1.79 | 0 |
| | *TUBA1B* | 1.78 | 0 |
| | *TOP2A* | 1.74 | 0 |
| | *STMN1* | 1.69 | 0 |
| | *MKI67* | 1.60 | 0 |
| | *UBE2S* | 1.59 | 0 |
| | *TIMP1* | 1.60 | 1.31E-134 |
| | *HIST1H4C* | 1.66 | 2.83E-130 |
| Sweat/Seba | *DCD* | 7.05 | 0 |
| | *SCGB2A2* | 5.49 | 0 |
| | *MUCL1* | 4.78 | 0 |
| | *SCGB1B2P* | 4.62 | 0 |
| | *SCGB1D2* | 4.23 | 0 |
| | *PIP* | 4.18 | 0 |
| | *AZGP1* | 3.17 | 0 |
| | *KRT19* | 2.50 | 0 |
| | *KRT7* | 2.00 | 0 |
| | *AQP5* | 1.95 | 0 |
| Schwann | *MPZ* | 2.59 | 0 |
| | *S100B* | 2.35 | 0 |
| | *GPM6B* | 1.99 | 0 |
| | *NRXN1* | 1.88 | 0 |
| | *PLP1* | 1.87 | 0 |

| | | | |
|---|---|---|---|
| | *CDH19* | 1.83 | 0 |
| | *VWA1* | 1.47 | 0 |
| | *CRYAB* | 1.87 | 4.26E-223 |
| | *PMP22* | 2.07 | 6.81E-198 |
| | *MBP* | 1.54 | 4.27E-23 |
| Plasma | *IGKC* | 7.48 | 0 |
| | *IGLC2* | 7.03 | 0 |
| | *IGHA1* | 6.25 | 0 |
| | *IGHG1* | 6.20 | 0 |
| | *IGLC3* | 5.78 | 0 |
| | *IGHG2* | 5.42 | 0 |
| | *IGHG3* | 5.16 | 0 |
| | *JCHAIN* | 4.31 | 0 |
| | *IGHG4* | 4.29 | 0 |
| | *IGLC7* | 4.27 | 0 |

**Supplementary Table 5: List of Top 10 genes overexpressed in the fibroblast sub-clusters.** Table showing top ten marker genes for each sub-cluster with average fold-change (FC) and adjusted p value. The fold change was calculated by comparing the expression profiles of target cluster with rest of the cluster.

| Cluster # | Gene | average logFC | adjusted p value |
|---|---|---|---|
| 0 | *APOE* | 1.67 | 0 |
| | *APOD* | 1.25 | 0 |
| | *GSN* | 1.17 | 0 |
| | *C7* | 1.13 | 0 |
| | *CFD* | 1.00 | 0 |
| | *MGP* | 0.98 | 0 |
| | *CXCL12* | 0.92 | 0 |
| | *MYOC* | 0.91 | 0 |
| | *CFH* | 0.88 | 0 |
| | *C3* | 0.79 | 0 |
| 1 | *WISP2* | 1.44 | 0 |
| | *PI16* | 1.19 | 0 |
| | *SLPI* | 1.01 | 0 |
| | *DCN* | 0.96 | 0 |
| | *FBLN1* | 0.95 | 0 |
| | *SFRP2* | 0.91 | 0 |
| | *CFD* | 0.88 | 0 |
| | *ANGPTL1* | 0.87 | 0 |
| | *WIF1* | 0.85 | 0 |
| | *TNXB* | 0.77 | 0 |
| 2 | *PTGDS* | 2.34 | 0 |
| | *CCL19* | 1.99 | 0 |
| | *APCDD1* | 0.90 | 0 |
| | *SFRP2* | 0.68 | 0 |
| | *PTN* | 0.60 | 0 |
| | *IL11RA* | 0.48 | 0 |
| | *APOE* | 0.77 | 1.4E-302 |
| | *CTSC* | 0.44 | 3E-239 |
| | *C3* | 0.65 | 4.1E-197 |

| | | | |
|---|---|---|---|
| | *CXCL12* | 0.48 | 2E-196 |
| 3 | *MMP1* | 2.77 | 0 |
| | *MMP3* | 2.70 | 0 |
| | *CHI3L1* | 2.57 | 0 |
| | *CHI3L2* | 2.20 | 0 |
| | *FTH1* | 1.87 | 0 |
| | *CXCL8* | 1.84 | 0 |
| | *MMP13* | 1.78 | 0 |
| | *CCL20* | 1.73 | 0 |
| | *MT2A* | 1.65 | 0 |
| | *CXCL1* | 1.65 | 0 |
| 4 | *POSTN* | 2.30 | 0 |
| | *ASPN* | 2.21 | 0 |
| | *TNC* | 1.78 | 0 |
| | *COL1A1* | 1.44 | 0 |
| | *COL12A1* | 1.43 | 0 |
| | *SFRP4* | 1.35 | 0 |
| | *TGFBI* | 1.34 | 0 |
| | *MMP11* | 1.32 | 0 |
| | *SPARC* | 1.30 | 0 |
| | *COL5A2* | 1.18 | 0 |
| 5 | *COMP* | 1.75 | 0 |
| | *APCDD1* | 1.42 | 0 |
| | *COL18A1* | 0.95 | 0 |
| | *LEPR* | 0.90 | 0 |
| | *CD9* | 0.85 | 0 |
| | *WIF1* | 0.83 | 0 |
| | *PCSK1N* | 0.81 | 0 |
| | *COL23A1* | 0.76 | 0 |
| | *F13A1* | 0.76 | 0 |
| | *THBS4* | 0.78 | 5E-244 |
| 6 | *PLA2G2A* | 2.69 | 0 |
| | *PRG4* | 1.92 | 0 |
| | *SFRP4* | 1.46 | 0 |
| | *HTRA3* | 1.40 | 0 |

| | | | |
|---|---|---|---|
| | *RARRES1* | 1.33 | 0 |
| | *IGFBP6* | 1.26 | 0 |
| | *MFAP5* | 1.19 | 0 |
| | *IGF1* | 1.03 | 0 |
| | *CD55* | 0.89 | 0 |
| | *FBN1* | 0.90 | 1.5E-257 |
| 7 | *NEAT1* | 1.34 | 0 |
| | *MT-ATP6* | 1.31 | 0 |
| | *MT-CO1* | 1.27 | 0 |
| | *MT-CO3* | 1.23 | 0 |
| | *MT-CO2* | 1.20 | 0 |
| | *MT-CYB* | 1.18 | 0 |
| | *MT-ND1* | 1.16 | 0 |
| | *XIST* | 1.13 | 0 |
| | *MT-ND5* | 1.12 | 0 |
| | *MT-ND4* | 1.07 | 0 |
| 8 | *GNB2L1* | 1.91 | 0 |
| | *ATP5E* | 1.70 | 0 |
| | *SELM* | 1.57 | 0 |
| | *TCEB2* | 1.31 | 0 |
| | *AC090498.1* | 1.29 | 0 |
| | *RPS17* | 1.25 | 0 |
| | *ATP5L* | 1.22 | 0 |
| | *SEPP1* | 1.19 | 0 |
| | *ALDOA* | 1.13 | 0 |
| | *MT2A* | 1.09 | 4.3E-159 |
| 9 | *NR2F2* | 1.62 | 0 |
| | *ANGPTL7* | 1.59 | 0 |
| | *C2orf40* | 1.33 | 0 |
| | *KLF5* | 1.29 | 0 |
| | *FOXD1* | 1.25 | 0 |
| | *TAGLN* | 1.24 | 0 |
| | *ITGA6* | 1.22 | 0 |
| | *TXNIP* | 1.19 | 0 |
| | *APOD* | 2.36 | 1.6E-262 |

| | | | |
|---|---|---|---|
| | *ID1* | 1.19 | 1.3E-181 |
| 10 | *IGFBP2* | 1.62 | 0 |
| | *CPE* | 1.45 | 0 |
| | *SFRP1* | 1.23 | 0 |
| | *FGFBP2* | 1.13 | 0 |
| | *OLFML2A* | 1.06 | 0 |
| | *RAMP1* | 0.78 | 0 |
| | *IGFBP5* | 1.29 | 2.1E-194 |
| | *SPRY1* | 0.75 | 2.1E-186 |
| | *PTN* | 0.86 | 6.4E-161 |
| | *TIMP3* | 0.96 | 2.1E-137 |
| 11 | *ACTA2* | 2.67 | 0 |
| | *TAGLN* | 2.17 | 0 |
| | *GNB2L1* | 1.92 | 0 |
| | *RGS5* | 1.61 | 0 |
| | *SELM* | 1.50 | 0 |
| | *NDUFA4L2* | 1.49 | 0 |
| | *C10orf10* | 1.47 | 0 |
| | *AC090498.1* | 1.41 | 0 |
| | *MYL9* | 1.55 | 6.8E-239 |
| | *SPARCL1* | 1.45 | 2E-234 |
| 12 | *MYH11* | 1.34 | 0 |
| | *RGS5* | 1.39 | 2.2E-306 |
| | *ACTA2* | 2.27 | 3.4E-275 |
| | *TAGLN* | 2.23 | 2.2E-230 |
| | *NR2F2* | 1.20 | 1.5E-190 |
| | *DEPP1* | 1.18 | 1.2E-143 |
| | *MYL9* | 1.26 | 4.7E-132 |
| | *TPM2* | 1.21 | 1.7E-116 |
| | *C11orf96* | 1.39 | 1.9E-104 |
| | *CCL19* | 1.15 | 2.38E-25 |
| 13 | *PTTG1* | 1.92 | 0 |
| | *TOP2A* | 1.89 | 0 |
| | *STMN1* | 1.85 | 0 |
| | *UBE2S* | 1.82 | 0 |

| | | | |
|---|---|---|---|
| | *CENPF* | 1.79 | 0 |
| | *TUBA1C* | 1.63 | 4.1E-226 |
| | *H2AFZ* | 2.00 | 1.3E-196 |
| | *TUBA1B* | 1.62 | 3.5E-161 |
| | *MMP1* | 1.72 | 4E-112 |
| | *MT2A* | 1.69 | 1.3E-101 |

#### Supplementary materials

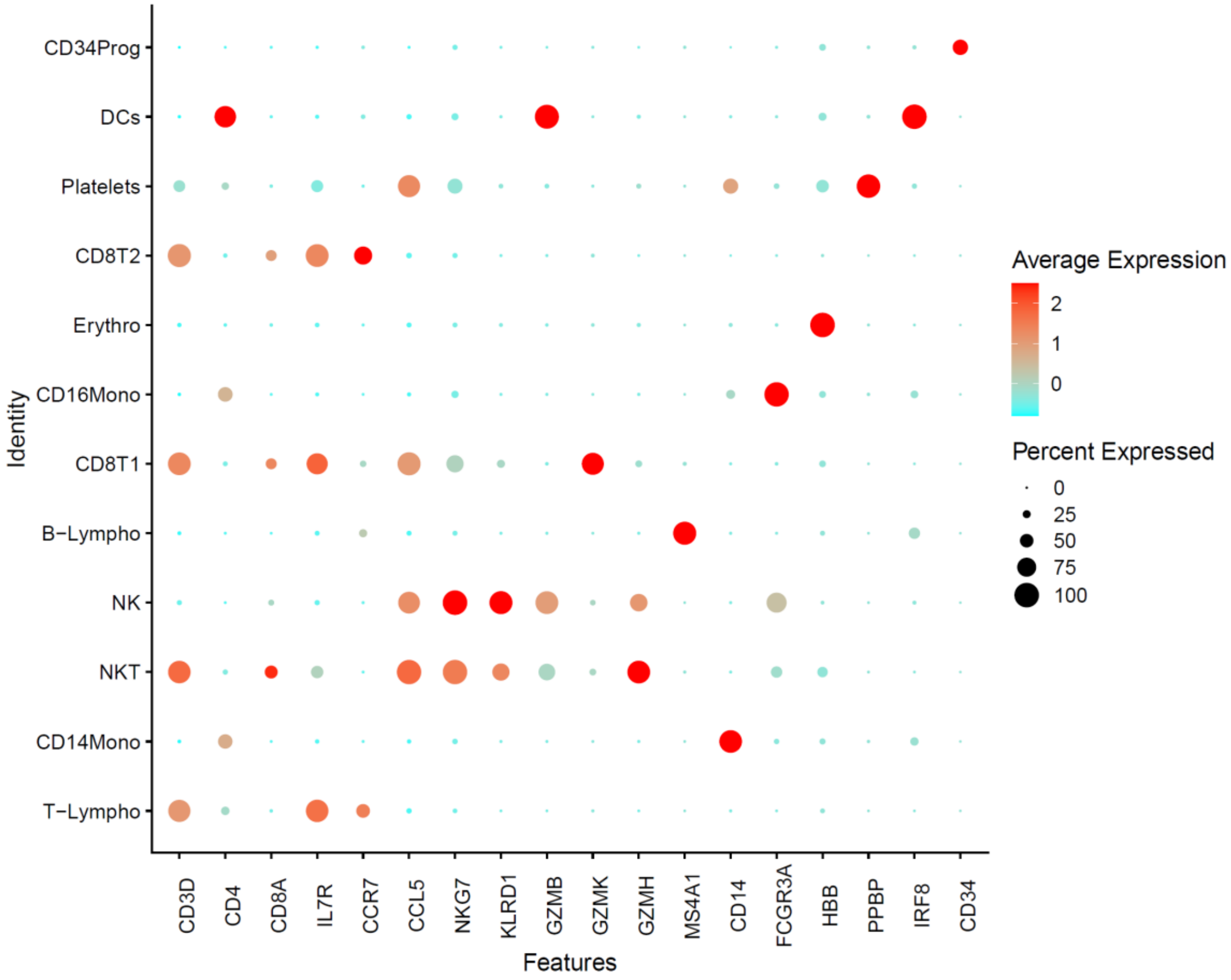


**Supplementary material 1: PBMC cell cluster annotation.** Cell annotation was performed based on expression of canonical marker genes. Dot plot showing expression of markers genes in 12 cell types. Size of dots indicates percentage of cells in each cluster that are expressing the marker gene; color represents averaged scaled expression levels; cyan: low, red: high.

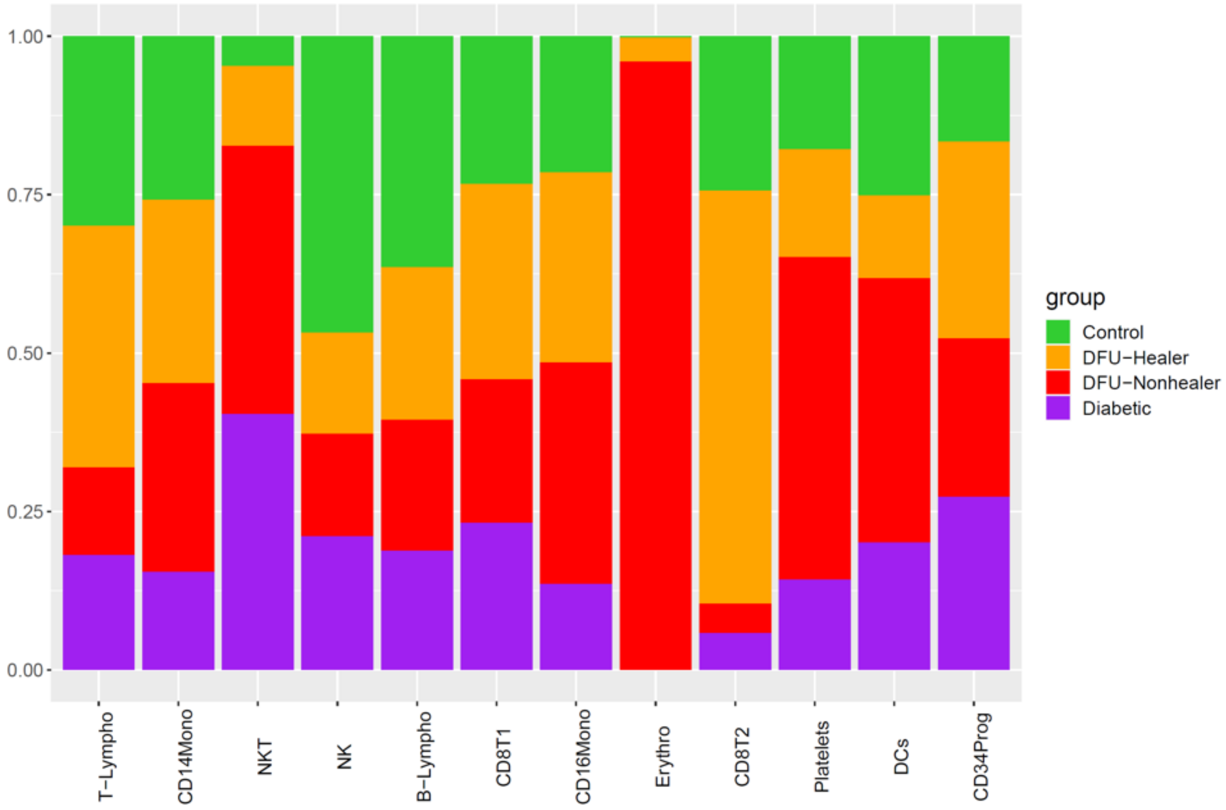


**Supplementary material 2: Distribution of cells from various clinical groups in PBMC cell types.** Stacked bar plots showing the proportions of different cell types across the different clinical groups (Green: Healthy controls, Orange: DFU-Healers, Red: DFU-Non-healers, Purple: non-DFU DM).

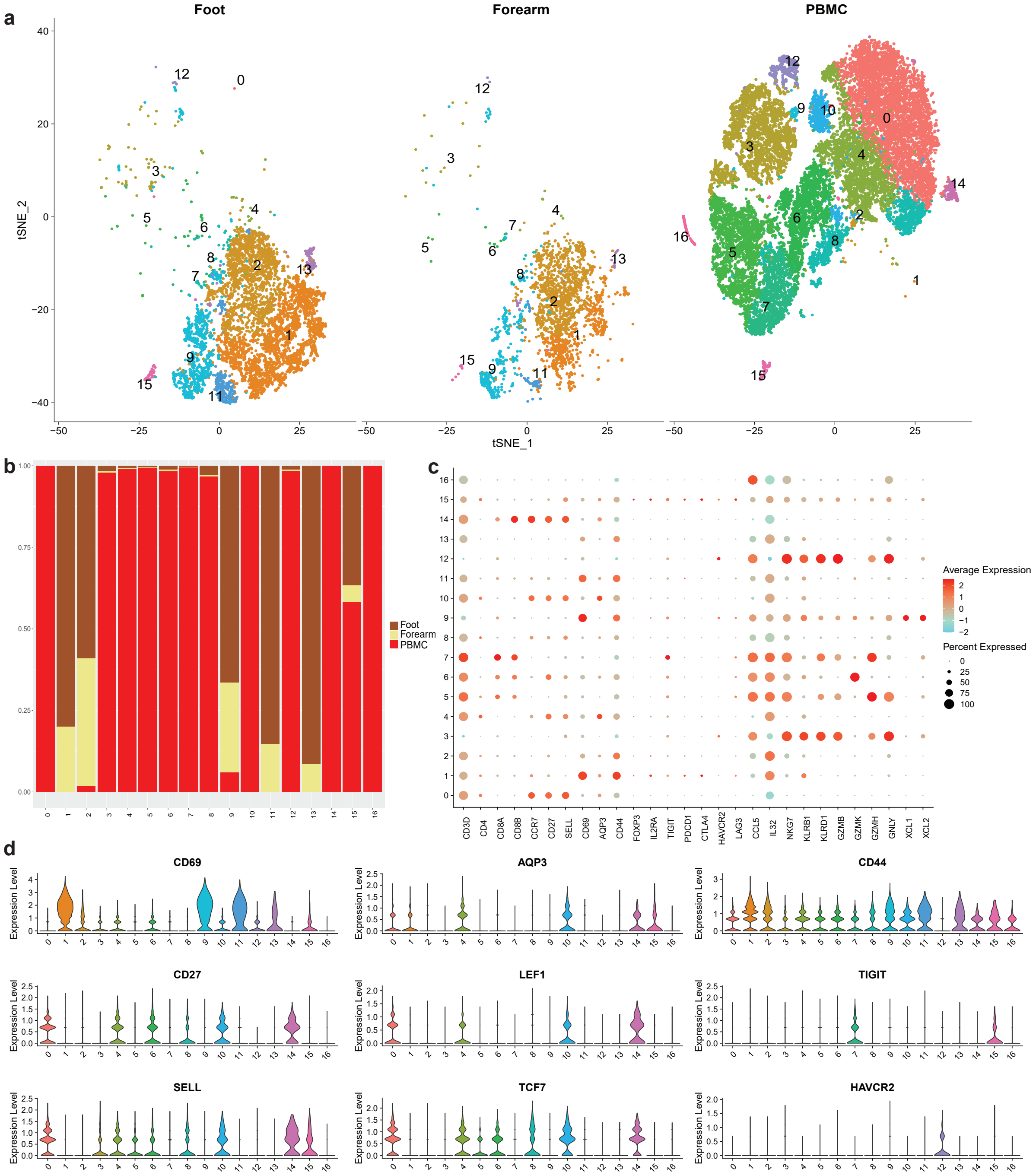


**Supplementary material 3: T cells and NK cells subset analysis.** (a) Split t-SNE plots based on anatomical sites, (b) Bar plot showing composition of each T/NK sub-cluster, (c) Dot plot showing expression of markers genes in different cell clusters for cell type annotation. Size of dots indicates percentage of cells in each cell cluster expressing the marker gene; color represents averaged scaled expression levels; cyan: low, red: high, (c) Violin plots showing expression of T cell marker genes for: activation (*CD69*$^+$, *CD44*$^+$), naive (*CD27*$^+$, *SELL*$^+$, *LEF1*$^+$), memory (*AQP3*$^+$), differentiation (*TCF7*$^+$) and exhaustion (TIGIT$^+$, HAVCR2$^+$).

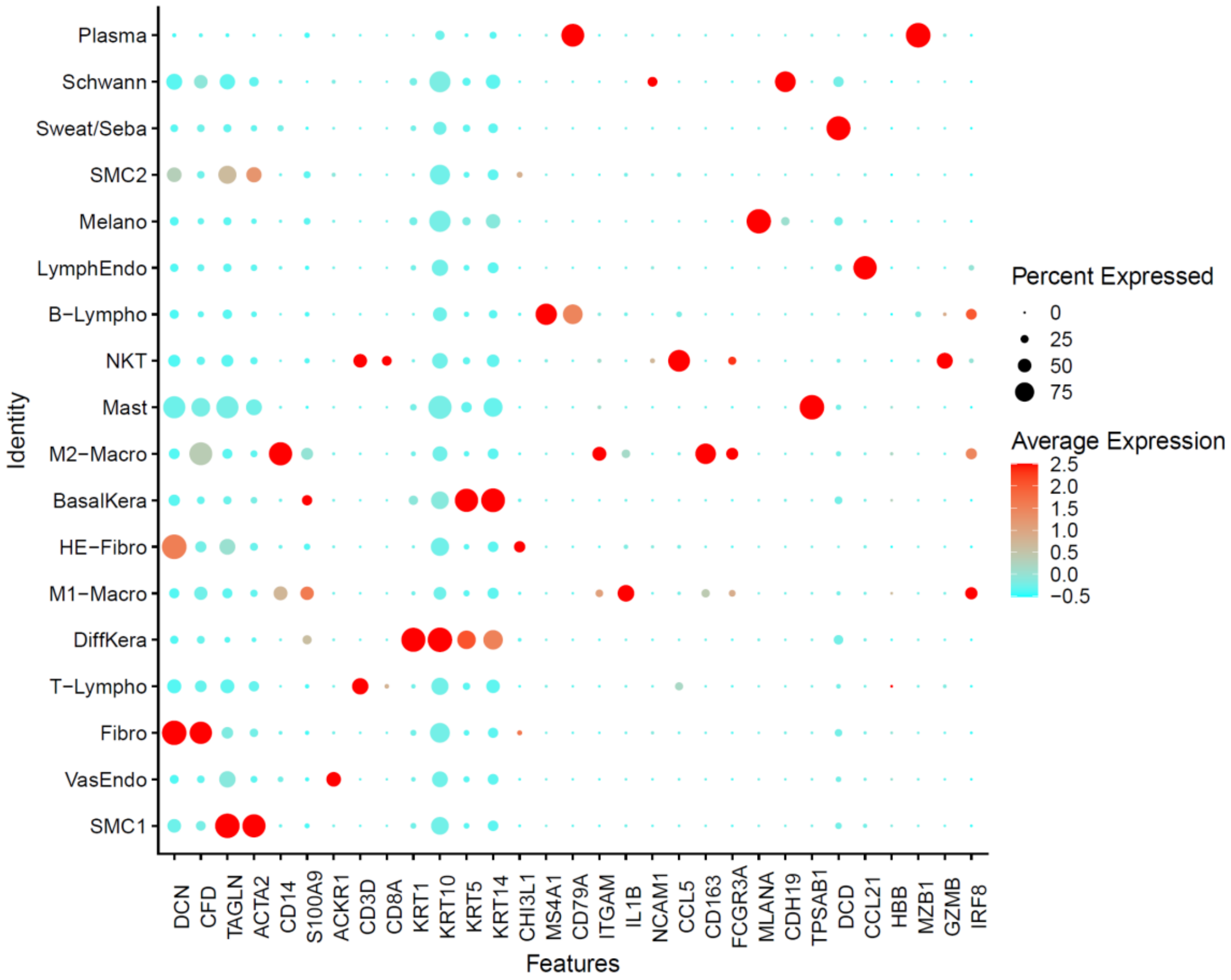


**Supplementary material 4: Annotation of foot clusters based on expression of marker genes.** Dot plot showing expression of markers genes in 18 clusters. Size of dots indicates percentage of cells in each cell cluster expressing the marker gene; color represents averaged scaled expression levels; cyan: low, red: high.

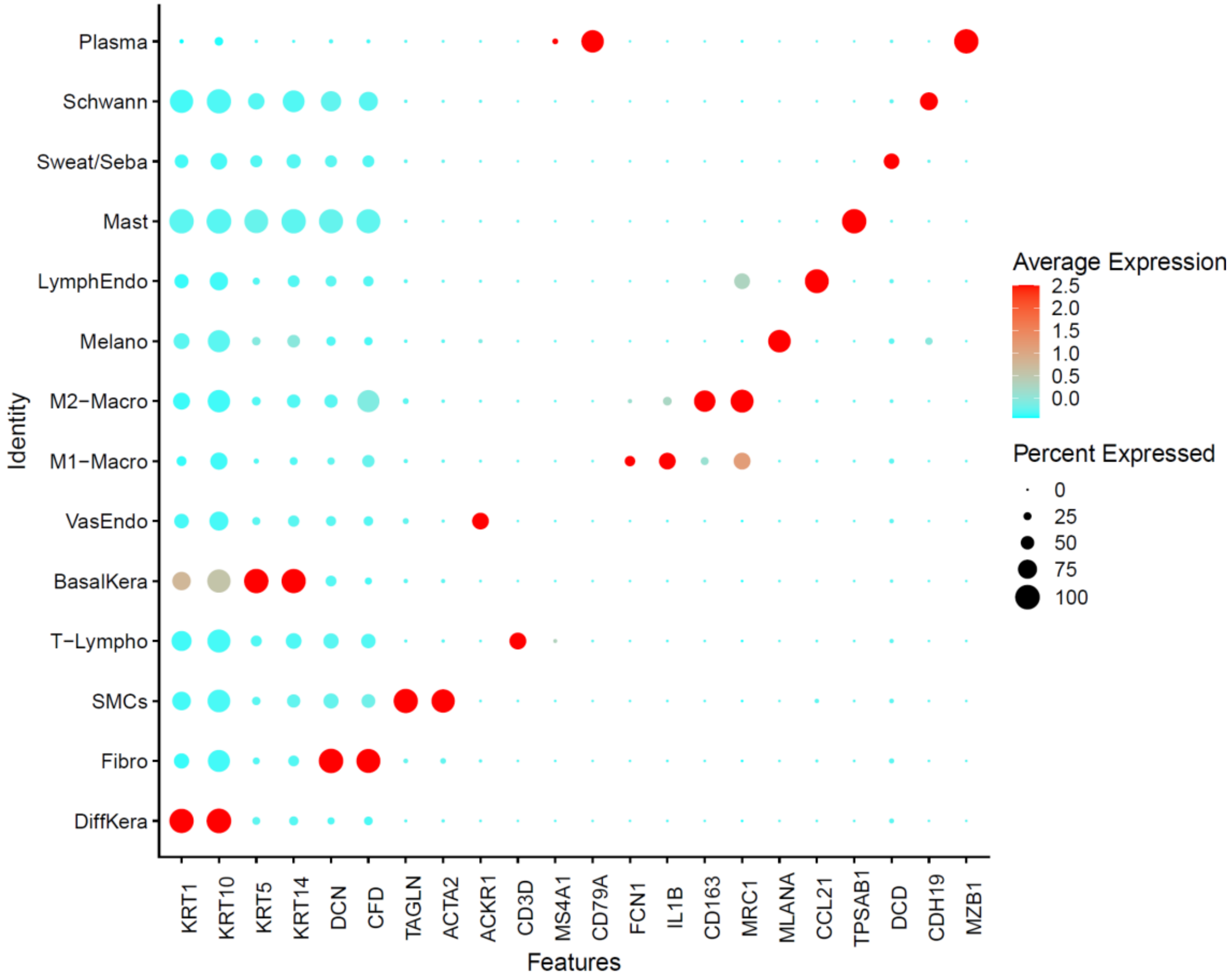


**Supplementary material 5: Cell type annotation of forearm cell clusters.** Dot plot showing expression of markers genes in 14 cell sub-types. Size of dots indicates percentage of cells in each cell cluster expressing the marker gene; color represents averaged scaled expression levels; cyan: low, red: high.

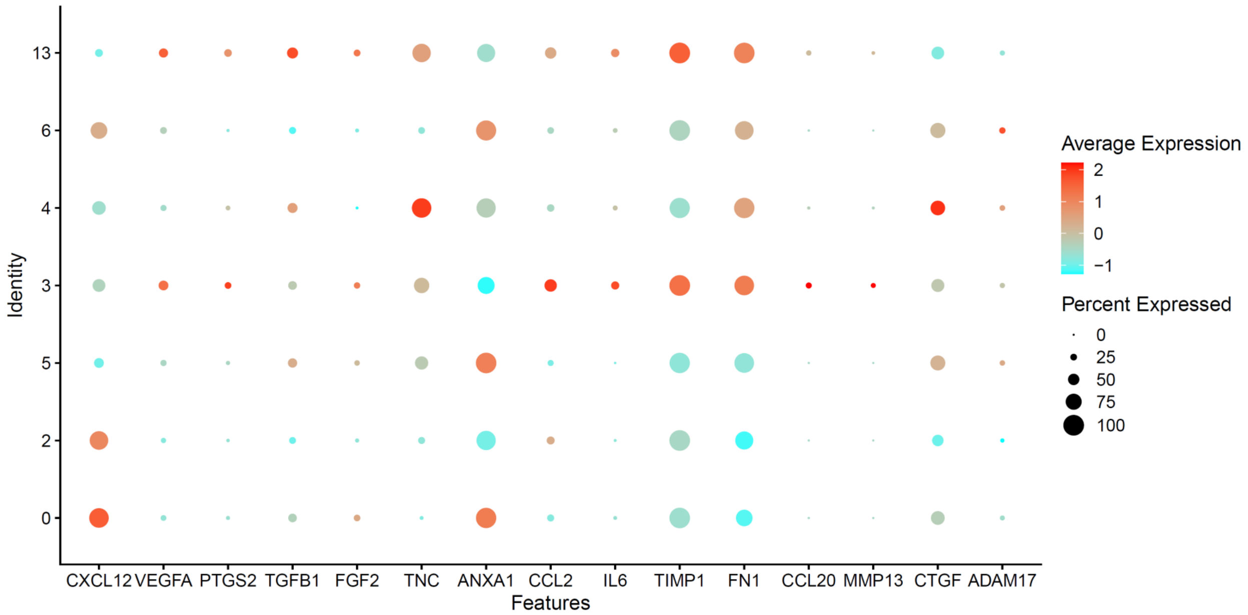


**Supplementary material 6.** Dot Plot shows the expression of some of the top ligands based on Pearson correlation expressed by the healer-enriched fibroblasts. X-axis shows the genes and Y-axis the cluster number. DFU-Healer enriched clusters (3, 4, 6, 13) show higher expression of *FN1, MMP13, TIMP1, CCL2,* IL6 and *CCL20* relative to DFU-Non-healer specific fibroblast clusters (0, 2, 5).

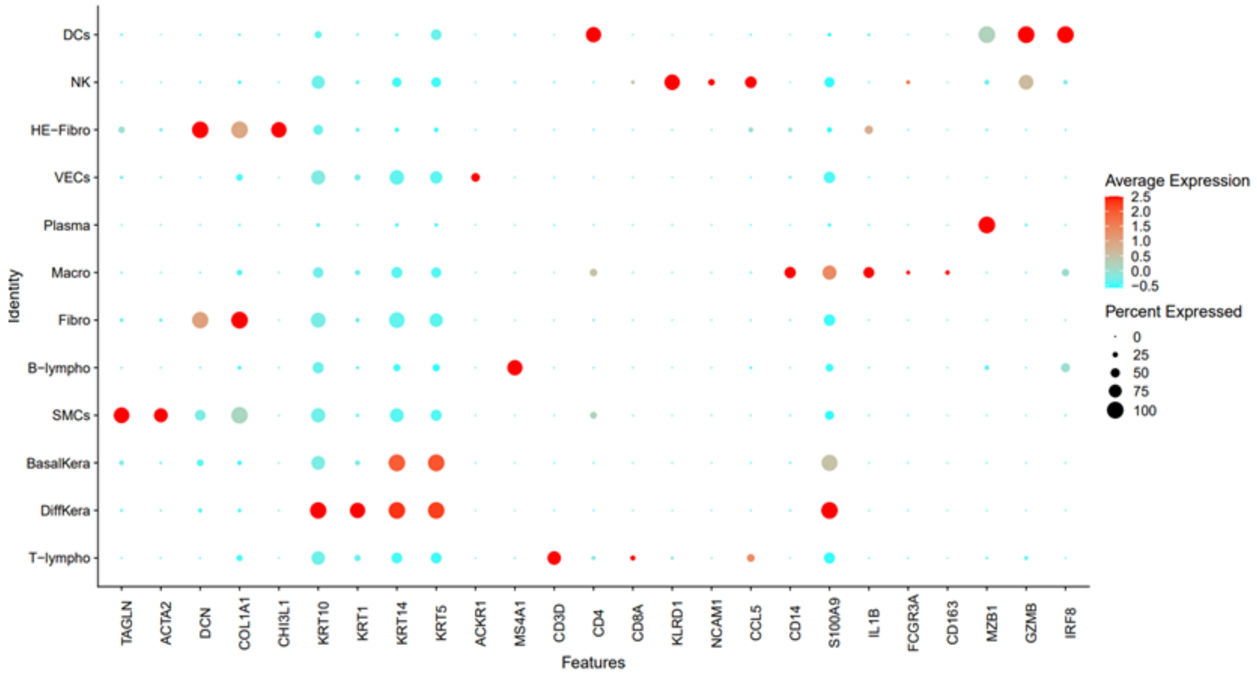


**Supplementary material 7: Cell type annotation of pressure sore cell clusters.** scRNA-Seq analysis was performed on skin specimens of the same patient from three different sites: wound bed, wound edge, and non-wound excess skin from a pressure sore excision. Dot plot showing expression of markers genes in the identified 12 cell types. X-axis shows the genes and Y-axis the cell type. Size of dots indicates percentage of cells in each cell cluster expressing the marker gene; color represents averaged scaled expression levels; cyan: low, red: high.
